# Phenotypic and genomic characterization of *Bacillus sensu lato* for their biofertilization effect and plant growth promotion features in soybean plants

**DOI:** 10.1101/2023.09.05.556405

**Authors:** P. Torres, N. Altier, E. Beyhaut, N. Martin, P. Fresia, S. Garaycochea, E. Abreo

## Abstract

*Bacillus sensu lato* were screened for their capacity to mineralize organic phosphorus (P) and promote plant growth, improving nitrogen (N) and phosphorus nutrition of soybean plants. Isolates were first identified based on their genomic sequences through TYGS and ANII. ILBB95, ILBB510 and ILBB592 were identified as *Priestia megaterium*, ILBB139 as *Bacillus wiedmannii*, ILBB44 as a member of a sister clade of *B. pumilus* (together with a human pathogenic strain), ILBB15 as *Peribacillus butanolivorans* and ILBB64 as *Lysinibacillus* sp. These strains were evaluated for their capacity to mineralize organic P as sodium phytate and solubilize inorganic P forms in liquid medium. These *in vitro* assays allowed the strains to be ranked according to their P mobilization potential, with ILBB15 and ILBB64 showing the highest orthophosphate production from phytate, ILBB592 the lowest and ILBB510 nil. In addition, features related to their rhizocompetence and plant growth promotion were evaluated *in vitro* and *in silico.* Finally, plant bioassays were deployed to assess the effect of the co-inoculation of *Bacillus s.l.* strains and rhizobial inoculant on nodulation, plant growth and nutrition. *In planta* bioassays showed that *B. pumilus* ILBB44 and *P. megaterium* ILBB95 increased P absorbed in plants grown on a poor substrate of sand and vermiculite and also on the richer mix of sand, vermiculite and peat. *Priestia megaterium* ILBB592 increased rhizobial nodulation and N content in plants grown on sand, vermiculite and peat mixture only. ILBB15 reduced plant growth and nutrition on both substrates. Genomes of ILBB95 and ILBB592 were characterized by genes related with plant growth and biofertilization whereas ILBB15 was differentiated by genes related to bioremediation. *Priestia megaterium* ILBB592 can be described as nodule-enhancing rhizobacteria (NER) and together with ILBB95, can be envisaged as prospective PGPR with the capacity to exert a positive effect on N and P nutrition of soybean plants.

## Introduction

The *Bradyrhizobium*-soybean symbiosis is considered one of the most important natural relations exploited in agriculture as it allows provision of atmospheric N to extensively produced crops, reducing the risks usually associated with chemical fertilization (Chang et al., 2015; Hungria et al., 2015; Colino et al 2015; Zeffa et al., 2020). Phosphorus (P) is the second most important macronutrient in plant nutrition and plays a crucial role in plant processes, such as photosynthesis, root elongation, signal transduction and nitrogen (N) fixation (Korir et al. 2017). Inadequate P levels restrict roots growth, the process of photosynthesis, the translocations of sugar and other functions, which directly or indirectly influence N fixation by legumes (Chen et al., 2023). In soils, inorganic and organic P are present mostly in unavailable forms for plants, whereas a small fraction is in solution and therefore bioavailable for plants. Organic P in the form of phytate and less complex molecules like nucleotides are generally estimated to contribute between 30 and 50% of total P, being most abundant in soils with high organic matter content (Mullen, 2005). Therefore, plants and microbes need to solubilize inorganic P and mineralize organic P to release orthophosphate ions to the soil solution where orthophosphate can then be absorbed. To do so, plants and microbes depend on the excretion of organic acids, phosphatases and phytases (Richardson et al., 2011). Fungi like *Penicillium billiae* but mostly Gram-negative bacteria like *Pseudomonas* spp. have been selected for their P mobilization capacity to be used as biofertilizers (Sanchez et al., 2016, Shen et al 2016). However, in spite of its many advantages, spore forming *Bacillus* have not been widely studied for their P mobilization capacity (Gomez and Uribe, 2021). Jorquera et al. (2008) characterized isolates of *Bacillus* for the phytase activity and found that some isolates were capable of mineralizing organic P and reported on genes coding for phytase in these isolates. Additionally to the P mineralization and solubilization potential, *Bacillus* and other bacterial genera have been pointed as responsible for improved nodulation of legumes and increased N fixation mostly through IAA and ACC desaminase production (Alemneh et al 2020; Flores Duarte et al 2022).

Uruguayan soils have variable levels of total P, but available P is usually insufficient for most crops (Garaycochea et al 2020). The strategy to lift this limitation has been the addition of phosphate fertilizer, mostly soluble superphosphate, which readily becomes fixed by the soil. Being a non-renewable and imported resource (Hernandez and Zamalvide, 1998), excessive fertilization can lead to water contamination through run-off and soil erosion.

In order to develop a *Bacillus* P-biofertilizer for soybean, a collection of *Bacillus sensu lato* with potential capacity to solubilize inorganic P and mineralize organic P in solid media was further characterized by quantitatively measuring the production of orthophosphate in liquid media. In addition, relevant features that are related to their rhizocompetence and plant growth promotion were evaluated *in vitro* and *in silico*. The effects of coinoculating the described *Bacillus* together with rhizobia on soybeans seeds, most notably the effect on nodule formation, P and N nutrition were evaluated and discussed in view of the phenotypic and genomic characterization of the *Bacillus* isolates.

## Materials and Methods

### Biological materials

Isolates of *Bacillus sensu lato* known to form a halo on solid medium supplemented with sodium phytate (ILBB15, ILBB44, ILBB64, ILBB95, ILBB592, ILBB139) and a strain that was negative for this feature (ILBB510) were retrieved from the INIA Las Brujas Bacteria collection (ILBB) and further identified and characterized. For *Bacillus*-rhizobia co-inoculation bioassays, two strains of *Bradyrizhobium elkanii* (U-1301 and U-1302) were included as rhizobial inoculants of soybean seeds cv. Nidera 5909.

### Species identification

#### DNA extraction and Whole-genome Sequencing

A single colony of each strain was incubated at 25°C for 24 h at 170 rpm in LB medium. Subsequently, Genomic DNA was extracted using DNeasy Blood & Tissue Kit (Qiagen, Austin, Texas, USA), as recommended by the manufacturer. DNA was checked for quality using a NanoDrop and subjected to gel electrophoresis (1% of agarose). Sequencing of whole genome was performed on Ilumina HiSeq4000 plataform (Illumina, CA, USA) by Beijin Genome Institute (Beijin, China) to generate paired-end 150 bp reads, obtained 5 M reads per sample with a quality score of Q20 > 98%. The quality of raw reads sequence was checked using FASTQC v 0.11.12 (http://www.bioinformatics.babraham.ac.uk/projects/fastqc/). Sequences were trimmed and adapter removed using Trimmomatic (v0.36) (Bolger et al., 2014). The SPAdes V.3.11.1 (Bankevich et al., 2012) was used to assemble draft genomes with the parameters: careful mode, Sspace, GapFiller – Assembly Improvement. The draft genomes were submitted to the National Center for Biotechnology Information (NCBI) GenBank as part of BioProject PRJNA745340 and were automatically annotated using Prokka.

The draft genome sequences were uploaded to the Type (Strain) Genome Server (TYGS), a free bioinformatics platform available at https://tygs.dsmz.de for a whole genome-based taxonomic analysis (Meier-Kolthoff & Göker, 2019). Average nucleotide identity (ANI) was calculated by the MUMmer algorithm (ANIm) with JSpecies software (Richter et al. 2016) for the 7 strains in relation with available genomes of the closest species indicated by TYGS.

### Quantification of mineralization and solubilization capacity in liquid media

Liquid Angle media prepared according to Maougal et al. 2014 (g/L glucose 9.9, KNO3 0.1, CaSO4.2H2O 0.69, MgSO4.7H2O 0.49, 500 ul/L of solution A (Thiamine 0,1 g/L), 500 ul/L of solution B (Ferric citrate 10 g/L), and 200 ul/L of solution C (H3BO3 2.82, CuSO4.5H2O 0.098, MnSO4.H2O 3.08, NaMoO4.2H2O 0.29, ZnSO4.7H2O

4.4) and supplemented with either Sodium Phytate (3.96 g/L) or FePO4 (5.25 g/L), AlPO4 (2.87 g/L) or Ca3(PO4)2 (3.65 g/L) was used for quantification of orthophosphate production by the selected isolates. Bacteria suspensions were prepared and used to inoculate beakers containing the different P supplemented media by duplicate (final OD600nm = 0.8). Orthophosphate concentration was determined by molybdenum blue spectrophotometric method at time 0, 24 and 48 hours since inoculation for Po and 0,1, 7 days for Pi (De Negris et al., 2017) twice for each beaker at each time (technical replicates). Broth pH variation was measured in a separate aliquot of the culture broths, at the same time. Orthophosphate concentration by volume and time was calculated and expressed as umol_P / L h. A blank comprising uninoculated broth of each described medium was included.

### Traits associated with rhizocompetence

#### Motility assay

Swimming and swarming motility were evaluated for each selected strain on a plate assay using LB medium at 0.3% and 0.6% agar concentrations. Aliquots of 10 ml (OD600nm =0.6) were inoculated on LB medium and incubated at 30° for 24 h. Bacterial swimming and swarming was determined as the diameter of colony after 24 h. The assay was repeated three times with five replications.

#### Biofilm formation

Biofilm formation was quantified as described by O’Toole and Kolter (1998) with modifications after Stephanovic et al. (2007). First, bacteria were grown overnight in liquid Luria-Bertani (LB) medium and cell suspensions were standardized at 0.06 absorbance units measured at 600 nm. The standardized bacteria suspensions were diluted 1:100 in fresh LB broth and the soybean seed exudate separately. To obtain soybean seed exudates, 100 surface-disinfected seeds of Nidera 5909 variety were transferred into a 500-mL Erlenmeyer flask containing 100 mL sterile distilled water and still incubated for 24 hours at room temperature (Barbour et al., 1991, Yaryura et al., 2008). The exudates were sterile filtered through a 0.22 u filter.

Five wells of sterile microtiter plates were inoculated with 200 μl of the diluted broth of each strain, the negative control contained only the corresponding culture medium or the soybean seed exudate. Microtiter plates were incubated at 30°C without shaking for 72 hr. Microtiter plates wells were washed (H20), dried and finally, 150 μl of crystal violet (2%) was added. The crystal violet that remained in the biofilm was resuspended in 150 μl of 95% ethanol for 30 min. To quantify biofilm accumulation, the OD570 of each well was measured using a microtiter plate reader. Biofilm formation was normalized with respect to bacterial growth to obtain the specific biofilm formation (SBF), which was calculated by the following formula, SBF = (B - NC)/BG, where B is the amount of CV bound to the cells attached to the surface of the wells, NC is the negative control, and BG is the OD600 of bacterial growth (Yaryura et al., 2008).

### Traits associated with plant growth promotion

#### Nitrogen fixation

Each strain was inoculated in 50 ml glass bottles containing NFb medium with the addition of NH4Cl as a unique N source. Plates were incubated at 28°C for 7 days, and bacterial growth or halo formation were observed as qualitative evidence of the atmospheric N fixation (Baldani et al., 2014). A strain of *Herbaspirillum huttiense* known for its N fixation was included as positive control (Punschke & Mayans, 2011).

#### ACC production

1-Aminocycopropane-1-carboxylic acid (ACC) deaminase production was determined in DF medium containing 0.1% glucose and 3 mM ACC as the sole N source (Koo et al., 2010). Growth of the bacterial colony after 72 h was considered as positive for ACC deaminase.

#### IAA production

IAA production was detected by the method described by Brick et al. (1991) with some modification. Briefly, aliquots of 200 ml of bacterial suspension of each strain (OD540nm =0.5) were inoculated into 50 ml of TY medium amended with L-tryptophan (5 mM). After 48h of incubation at 150 rpm and 28°C, cells were harvested by centrifugation (10000 rpm for 10 min) and 250 ml of the supernatant was treated with 1 ml of Salkowski reagent (50 ml of perchloric acid (35%) and 1 ml of FeCl3 (0.5M)) for 30 min, under light protection. IAA was measured by measuring optical density at 530 nm. Commercial IAA as used as standard to quantify the IAA. The assay was repeated three times with five replications.

#### Hydrogen cyanamide production

Qualitative determination of hydrogen cyanamide (HCN) was performed according to Egan et al., (1998), where bacterial strains were streaked on 10% Tryptone Soy Agar (TSA) supplemented with glycine (4.4 g/ L). An autoclaved filter paper soaked with picric acid (0.5%) and Na2CO3 (2%) solution was fixed to the underside of the lids of petri plates and incubated for 48 h at 30°C. A change in colour from yellow to orange-brown was the indication of HCN production. The assay was repeated three times with five replications.

#### In planta bioassays

Three different experiments -which shared the seed inoculation procedure- were conducted under greenhouse conditions at the INIA Las Brujas, Uruguay, to assess nodulation and P and N nutrition. Seeds were first inoculated with a bacterial suspension of strains U-1301 and U-1302 of *Bradyrizhobium elkanii* (Active-N, concentration1x10^9^ CFU/mL) in plastic bags (150 ml/50 kg of seeds) and then planted in pots. Each pot was sown with 4 seeds, assigning ten pots to each treatment. Treatments consisted in seeds inoculated with *Bradyrizhobium elkanii* only (control) and seeds inoculated with *Bradyrizhobium elkanii* supplemented with a spore suspension of each strain of *Bacillus s.l.* separately.

For spore production, *Bacillus* strains were incubated in 2X SG medium (Leighton and Doi, 1971) at 28°C for 48 hrs and 180 rpm in a rotary shaker when sporulation was evident. For preparation of spore suspensions, spores were recovered by centrifugation (5000 rpm x 5 min), and the pellets were washed twice with distilled water. Finally, spore concentrations were adjusted to 1x10^6^ CFU ml^-1^ and stored at 4 C. Spore viability was checked before seed inoculation by plating a dilution and counting cfu/ml. Soybean seeds, already sown, were then inoculated with 1 mL of each *Bacillus* spore suspension by directly discharging this volume with a pipette onto the seeds.

After of 15 days from emergence, two plants were removed from each pot. Plants were watered with 50 ml of water every 3 days and fertilized with 10 ml of nutrient solution containing 83,0 nM sodium phytate without nitrogen, once a week.

#### Nodulation assessment

For nodulation assessment, pots were filled with a 1:1 substrate mixture of sand and vermiculite and seeded as previously described. Pots were kept in a greenhouse (25°C and 16 hours light/8 hours dark) for 30 days. After that period, roots were collected and number and weight of nodules in principal and secondary roots were recorded. Nodules were removed and placed in an oven at 65°C until constant dry weight was obtained (DW). DW was expressed as grams per plants (g/plant). This experiment was conducted twice.

#### P and N nutrition

Two independent trials were performed in two substrate mixes, with different content of organic matter. Pots were kept in greenhouse (28°C and 16 hours light/8 hours dark) for two months. After this period, plants were harvested to evaluate the effect of each treatment on plant nutrition. N and P were measured in the whole aerial part of plants (shoot and leaf together), with two plants per pots as experimental unit. Samples were dried in an oven at 65°C until constant dry weight (DW) was obtained. N and P concentration were expressed on per plant basis.

#### Genomic analysis

The bacterial genomes were analysed for the presence of plant growth promotion attributes by uploading the genome to the PLaBAse web server (Patz et al. 2021; Ashrafi et al. 2022). The KEGG annotations of the proteins of all strains were parsed into an IMG- like KEGG annotation file format via an in-house script and mapped against the plant growth promotion traits ontology with the PGPT-Pred tool, available on the web platform for plant-associated bacteria, PLaBAse (http://plabase.informatik.uni-tuebingen.de/pb/plabase.php) (Patz et al., 2021)

AntiSMASH 5.0 software (Blin et al. 2019) was used to identify the presence of gene clusters related to secondary metabolites. Specific genes related with P mobilization and nodulation were surveyed through a textual search. For this, the relationship of coding sequences (CDS) belonging to genes known for their activity in relation with inorganic phosphorus solubilization (ppqA, pqqB, pqqC, pqqD, ppx and ppa), organic phosphorus mineralization (phoX, phoA, phoD, olpA and phytases), and the P-starvation response regulation (phoR, phoB, phoP) were searched in each genome. In addition, the presence of genes coding for β-glucosidases (bglA, bglC, bglH and gmuD) that act on the plant isoflavone glycosides to start signalling between the plant and the rhizobia were searched in each draft genome. Protein domain assignment was performed using InterproScan.

### Statistical analysis

The normality and heteroscedasticity of the data were evaluated by shapiro-wilk. Each experiment was independently analysed by analysis of variance (ANOVA), and LSD multiple comparison test.

## Results

### Species identification

Draft genomes (Table 1) of the seven strains were initially submitted to TYGS for phylogenomic species identification (Supp. Figure 1). ILBB95, ILBB510 and ILBB592 clustered together with *Priestia megaterium* type strain ATCC 14581 (*formerly Bacillus megaterium*). *Priestia aryabhattai* JCM 13839 -type strain of the species- was also clustered with the *P. megaterium* clade although it was marked as a different species. ILBB15 clustered with *Peribacillus butanolivorans,* ILBB44 clustered in a sister clade with *B. pumilus* strain Bonn, next to a clade formed by *Bacillus pumilus* ATCC 7061 and NCTC 10337 type strains. ILBB139 clustered with *Bacillus wiedmannii* FSL W8-0169 and ILBB64 clustered with *Lysinibacillus xylanilyticus* DSM 23439, although they were labelled as different species. ANIm of strains in the *P. megaterium* clade (Table 1) ranged from 96.99 to 97.54 regarding *P. megaterium* type strain, whereas values were below 96 regarding *P. aryabhattai* type strain. ANIm of ILBB15 had a value of 97.76 with *Pe. butanolivorans*. ANIm of *B. pumilus* ILBB44 was 96.20 regarding the type strain of the species and 98.61 regarding the Bonn strain of *B. pumilus.* ANIm value of 96.43 of ILBB139 supported the TYGS phylogenomic association with *B. wiedmannii* whereas ILBB64 had an ANIm of 91.63 with the type strain of the species *Ly. xylanilyticus*.

**Table 1.** Draft genome data of *Priestia* and *Bacillus* strains and ANIm values with related strains indicated in the TYGS phylogenomic tree.

| Strain | Size | Contigs | GC [%] | ANIm to megaterium ATCC 14581 T | ANIm to aryabhattai B8W22 = JCM 13839 T | ANIm B. pumilus NCTC 10337 T | ANIm B. pumilus Bonn | ANIm B. wiedmannii FSL W8-0169 T | ANIm Pe. butanolivorans DSM 18926 T | ANIm Ly. xylanilyticus DSM 23493 T |
| --- | --- | --- | --- | --- | --- | --- | --- | --- | --- | --- |
| ILBB592 | 5,402,295 | 19 | 37.8 | 96.99 | 95.64 |  |  |  |  |  |
| ILBB510 | 5,600,674 | 90 | 37.7 | 97.54 | 95.66 |  |  |  |  |  |
| ILBB95 | 5,392,433 | 35 | 37.7 | 97.50 | 95.63 |  |  |  |  |  |
| ILBB 44 | 3,642,418 | 15 | 41.5 |  |  | 96.20 | 98.61 |  |  |  |
| ILBB15 | 6,093,790 | 145 | 37.7 |  |  |  |  |  | 97.76 |  |
| ILBB139 | 5,601,619 | 23 | 35.1 |  |  |  |  | 96.43 |  |  |
| ILBB64 | 5,126,053 | 45 | 36.9 |  |  |  |  |  |  | 91.63 |

### Strain characterization

*Bacillus* strains were evaluated for characteristics associated with rhizocompetence (biofilm formation, swimming, swarming) and growth promotion (IAA, siderophores, ACC deaminase) (Table 2). Swimming motility, an individual cell movement phenomenon dependent on the polar flagella rotation, was observed in all the strains but was relatively higher in two of them (ILBB95, ILBB44), intermediate in three of them (ILBB592, ILBB64 and ILBB139) and relatively lower in ILBB510 and ILBB15. Among these, swarming motility was high also in ILBB64 and ILBB44. Regarding indole-3- acetic acid (IAA), five strains produced IAA (ranging from 2 to 49 mg/ml), with ILBB64 showing the highest levels. Only two strains were negative for IAA production (ILBB15 and ILBB44). Nitrogen fixation determined by bacterial growth in NFb medium (nitrogen free), was observed in three strains (ILBB95, ILBB139, ILBB510). Among seven strains, four showed ACC deaminase activity (ILBB95, ILBB44, ILBB592, ILBB510), and three strains could produce HCN (ILBB15, ILBB64, ILBB592). Biofilm formation was distinctly high in ILBB592 and ILBB64 when grown on seed extract, whereas ILBB15 biofilm was higher on LB medium and lower on seed extract.

**Table 2.** *In vitro* P mobilization, rhizocompetence and PGPR traits displayed by the bacterial strains.

| Strain | P Org<br>mineralization | P Inorg<br>solubilization | ACC | Nitrogen | IAA | HCN | SBF |  | swimming | swarming |
| --- | --- | --- | --- | --- | --- | --- | --- | --- | --- | --- |
|  | (P-umol/L) | (P-umol/L) | Desaminase | Fixation | (mg/ml) |  | LB | Seed Ext. - | (mm) | (mm) |
| **ILBB15** | 7 | 95 | - | - | - | + | 2.3 ± 0.2 | 0.6 ± 0.2 | 33,7 ± 2,3 | 17,2 ± 6,1 |
| **ILBB64** | 5.7 | 120 | - | - | 49 ± 1 | + | 0.2 ± 0.02 | 1.3 ± 0.23 | 42 ± 8,7 | 85 ± 0 |
| **ILBB95** | 3.8 | 60 | + | + | 11 ± 0,4 | - | 0.2 ± 0.05 | 0.7 ± 0.8 | 85 ± 0 | 12,7 ± 1,6 |
| **ILBB139** | 3.7 | 125 | - | + | 3 ± 0,6 | - | 0.4 ± 0.09 | 0.5 ± 0.04 | 41,3 ± 1,4 | 13,5 ± 3,5 |
| **ILBB44** | 3.7 | 105 | + | - | - | - | 0.2 ± 0.02 | 0.3 ± 0.05 | 71 ± 6,1 | 59,3 ± 11 |
| **ILBB592** | 2.7 | 80 | + | - | 8 ± 1 | + | 0.9 ± 0.19 | 3.9 ± 0.23 | 44,7 ± 1,2 | 11 ± 0,4 |
| **ILBB510** | 0.0 | 0.0 | + | + | 2 ± 0,1 | - | 0.4 ± 0.08 | 0.4 ± 0.08 | 37 ± 3,5 | 10,7 ± 0,6 |

Positive (+): Growth of bacterial and/or colour change of the medium, negative (-): No growth of bacterial on medium.

**Table 3.** Presence/absence of selected genes involved in solubilization of inorganic P, mineralization of organic P, and P starvation response in Bacillus *sensu lato* strains.

|  | Inorganic P solubilization |  |  |  |  |  | Organic P mineralization |  |  |  |  |  | P-starvation response regulation |  |  |
| --- | --- | --- | --- | --- | --- | --- | --- | --- | --- | --- | --- | --- | --- | --- | --- |
|  | Pyrroloquinoline quinone synthase |  |  |  | Exopolyphosphatase | Pyrophosphatase | Alkaline phosphatase |  |  | Non specific acid phosphatase |  | PTP-like |  |  |  |
|  | pqq A | pqq B | pqq C | pqq D | ppx | Ppa | ph oX | ph oA | ph oD | olpA | ywjP | phytase | phoR | phoB | phoP |
| ILBB 592 | 0 | 1 | 1 | 0 | 1 | 1 | 1 | 1 | 1 | 1 | 1 | 0 | 1 | 1 | 1 |
| ILBB 510 | 0 | 1 | 1 | 1 | 1 | 1 | 1 | 1 | 1 | 1 | 1 | 0 | 1 | 1 | 1 |
| ILBB 95 | 0 | 1 | 1 | 0 | 1 | 1 | 1 | 1 | 1 | 1 | 1 | 0 | 1 | 1 | 1 |
| ILBB 44 | 0 | 1 | 1 | 1 | 1 | 1 | 0 | 1 | 0 | 1 | 1 | 0 | 1 | 0 | 1 |
| ILBB 139 | 0 | 1 | 1 | 0 | 1 | 1 | 0 | 1 | 0 | 1 | 1 | 0 | 1 | 0 | 1 |
| ILBB 64 | 0 | 1 | 0 | 0 | 1 | 1 | 1 | 1 | 0 | 1 | 0 | 1 | 1 | 1 | 1 |
| ILBB 15 | 0 | 1 | 1 | 1 | 1 | 1 | 1 | 1 | 1 | 1 | 1 | 0 | 1 | 1 | 1 |

**Table 4.** Presence of β-glucosidase genes involved in signalling conducive to nodulation.

| Strain | β-glucosidase genes | | | |
| --- | --- | --- | --- | --- |
|  | bglA | bglC | bglH | gmuD |
| ILBB15 | 0 | 1 | 0 | 0 |
| ILBB44 | 1 | 1 | 1 | 1 |
| ILBB64 | 0 | 0 | 0 | 0 |
| ILBB139 | 0 | 1 | 0 | 0 |
| ILBB95 | 1 | 0 | 1 | 0 |
| ILBB592 | 1 | 1 | 1 | 0 |
| ILBB510 | 1 | 0 | 1 | 0 |

### Effect of co-inoculation on nodulation

Seven strains characterized in the laboratory were included in plant bioassays performed under greenhouse conditions to evaluate the effect of the co-inoculation of soybeans seeds with *Bacillus* strains and *Bradyrhizobium elkanii* on nodulation. Thirty days after sowing, it was found that the nodular biomass expressed as nodule dry weight was 10.5 % and 21.1% higher (P>0.05) than the control treatment (only inoculated with *Bradyrhizobium elkanii*) when soybeans seeds were co-inoculated with strains ILBB95 and ILBB592, respectively. An opposite effect was observed on nodular biomass when seeds were co-inoculated with the strain ILBB15, which significantly reduced (P>0.05) nodular biomass compared to the control treatment.

### Effects of co-inoculation on P and N content

In the bioassays to evaluate the effect on P and N nutrition (60 days), the selected strains were evaluated in two types of substrates, which differed in the content of organic matter: sand-vermiculite substrate (SV) and sand-peat-vermiculite substrate (SPV). In both bioassays ILBB44 and ILBB95 strains presented the highest concentration of P per plant, and these results were statistically different (P>0.05) with an increase of 24% and 31%, respectively, compared to the control treatment, when evaluated in the poor sand-vermiculite substrate without the addition of peat (Fig 2a and 3a). Regarding N content (Figure 2b and 3b), the co-inoculation of the *Bacillus* strains showed statistically significant increases with respect to the control treatment (only inoculated with *Bradyrhizobium elkanii)* when the soybean seeds were co-inoculated with strains ILBB44, ILBB592, ILBB139 and ILBB510 in the SPV substrate.

**Figure 1.**
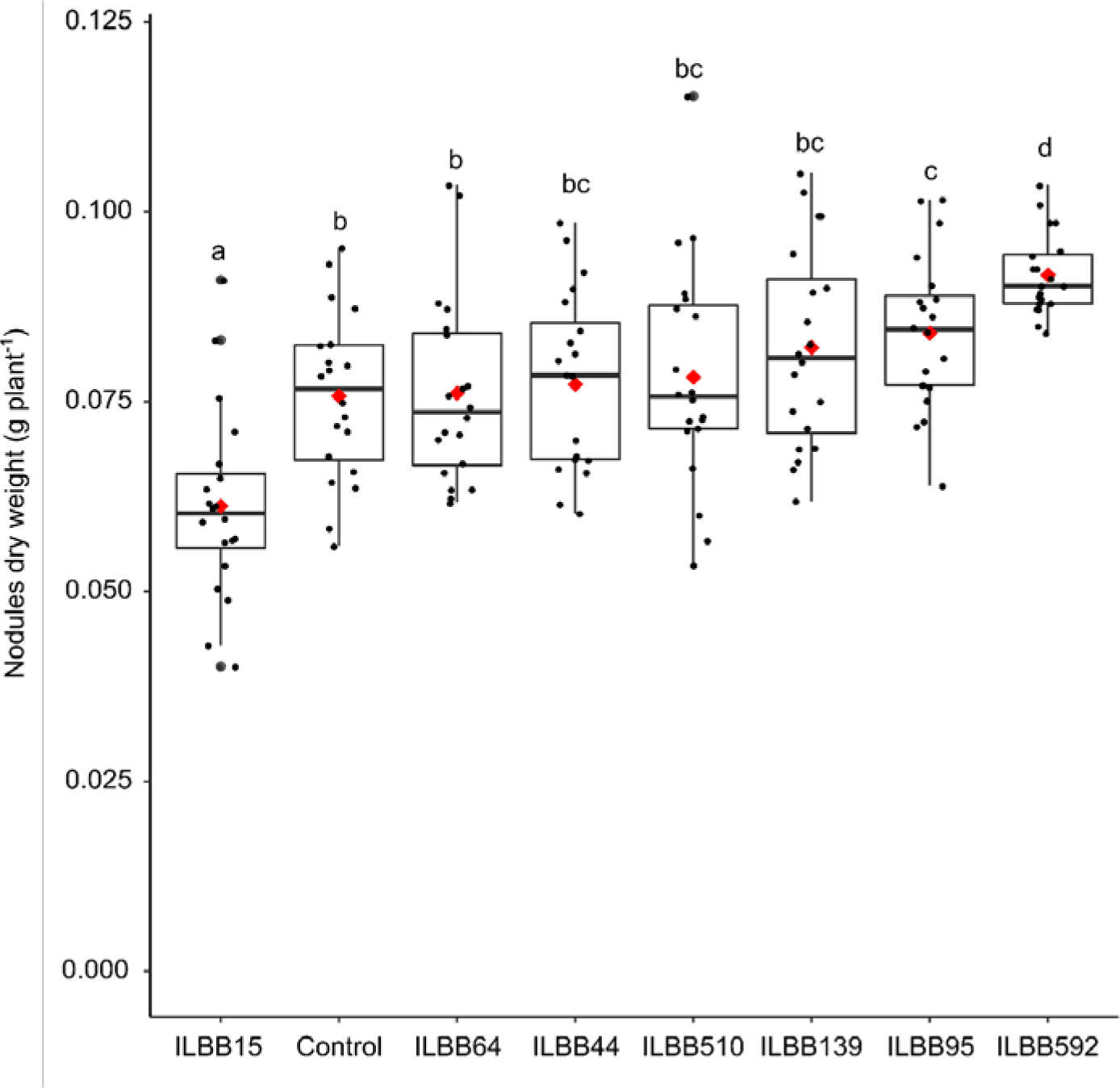
Effect of Co-inoculation of *Bradyrhizobium elkanii* with seven *Bacillus s.l*. strains on nodule dry weight in soybean plants grown on sand vermiculite mix under greenhouse conditions. Control: seeds inoculated with *Bradyrizhobium elkanii* only. Boxes contain observed values and mean of 2 repetitions and 10 replicates. Statistically different means are indicated by different letters (MLGM).

**Figure 2.**
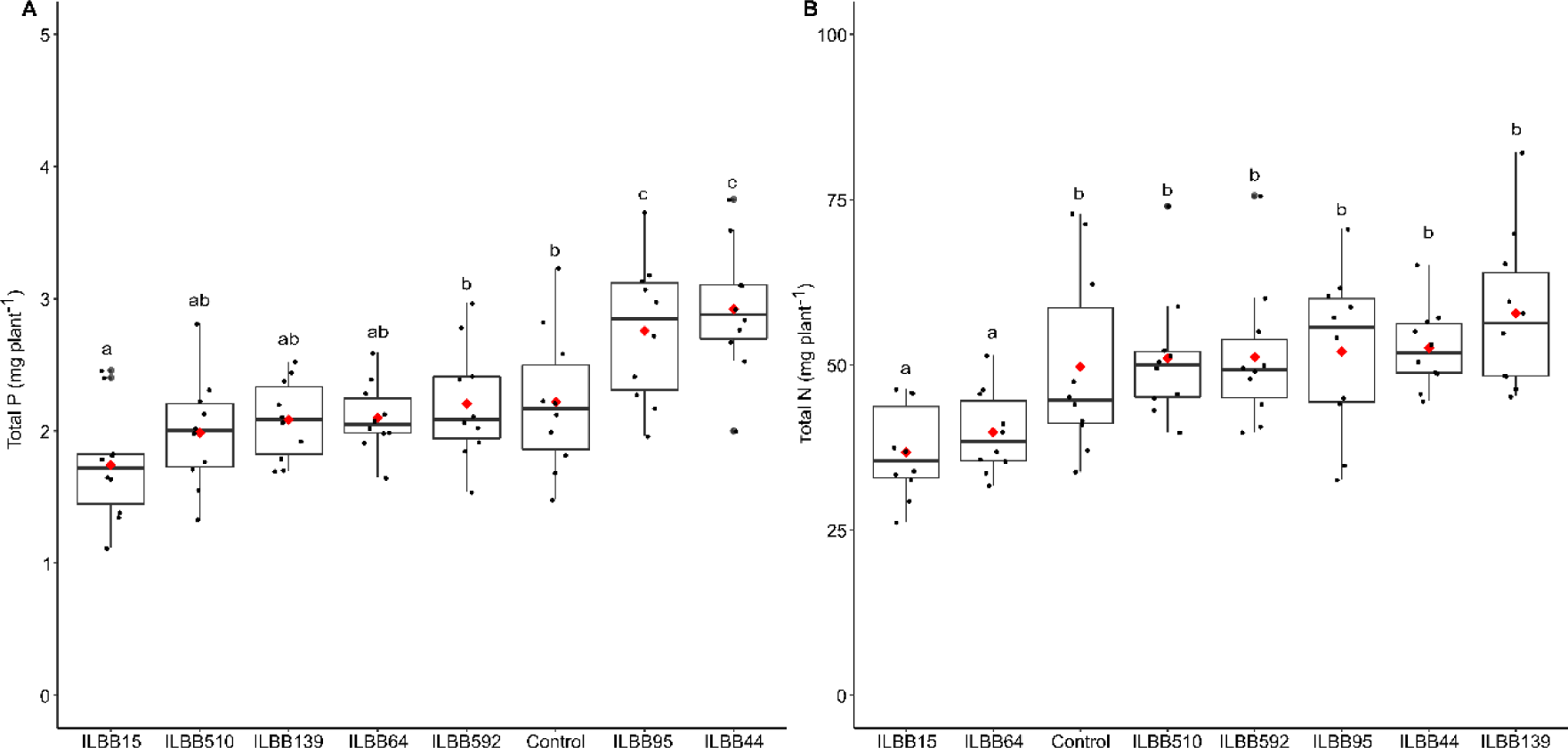
Effect of Co-inoculation of *Bradyrizhobium elkanii* with seven *Bacillus s.l.* strains on Phosphorus (A) and Nitrogen (B) content of plants grown on sand vermiculite mix under greenhouse conditions. Control: seeds inoculated with *Bradyrizhobium elkanii* only. Boxes contain observed values and mean of 10 replicates. Statistically different means are indicated by different letters (Fisher LSD).

**Figure 3.**
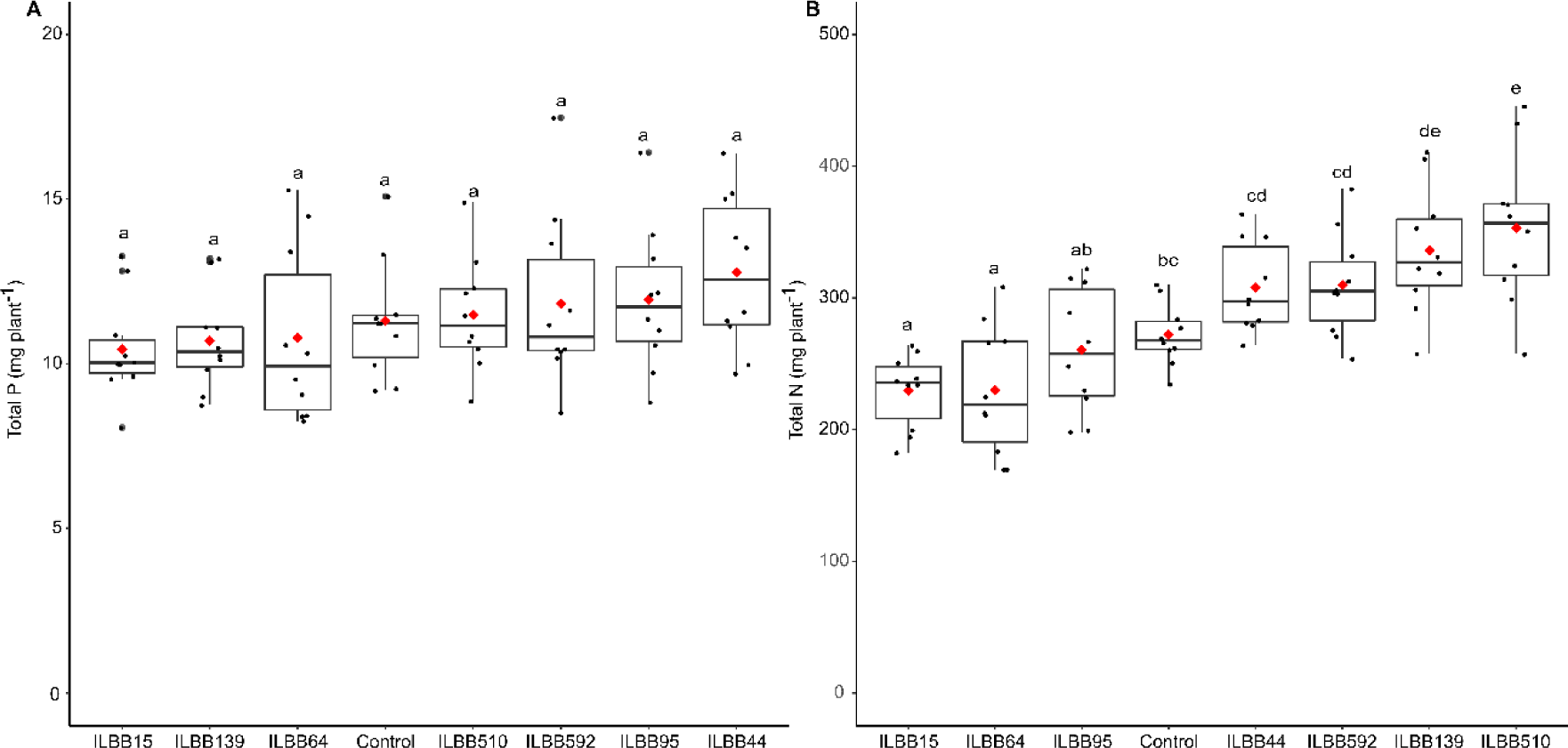
Effect of Co-inoculation of *Bradyrizhobium elkanii* with seven *Bacillus sensu lato* strains on Phosphorus (A) and Nitrogen (B) content on soybean plants grown in pots containing sand-peat-vermiculite mix under greenhouse conditions. Control: seeds inoculated with *Bradyrizhobium elkanii* only. . Boxes contain observed values and mean of 10 replicates. Statistically different means are indicated by different letters (Fisher LSD).

The co-inoculation with the strain ILBB15 in both substrates had a negative effect on P and N contents of soybean plants, compared to the control treatment. In addition, it was observed that the co-inoculation with the strain ILBB64 significantly reduced (P<0.05) the N content of soybean plants in both substrates (SV and SPV).

### *In Silico* Analysis of the Genome for Plant Growth Promoting Traits Functional PGPT annotation

An *in silico* search for plant growth-promoting genes was carried out in the draft genomes of the seven strains used in bioassays (Figure 4, 5). We found that strains identified as *Priestia megaterium* (ILBB592, ILBB95 and ILBB510), showed a higher number of genes encoding proteins with direct effects on plants, related to biofertilization (N acquisition, P and K solubilization) and phytohormone synthesis. These strains also had higher number of genes that encode traits that aid in plant system colonization, stress control-biocontrol and neutralizing abiotic stress, principally genes involved in neutralizing salinity stress (indirect effect). The strains ILBB15 (*Peribacillus butonolivorans*), which in general had a negative effect on soybean plants, had more genes related with resistance to heavy metals such as zinc, iron, nickel and also more genes involved in xenobiotic biodegradation. Also, the strains ILBB15, ILBB139, ILBB64 and ILBB44 were characterized by having more genes ascribed to competitive exclusion.

**Figure 4.**
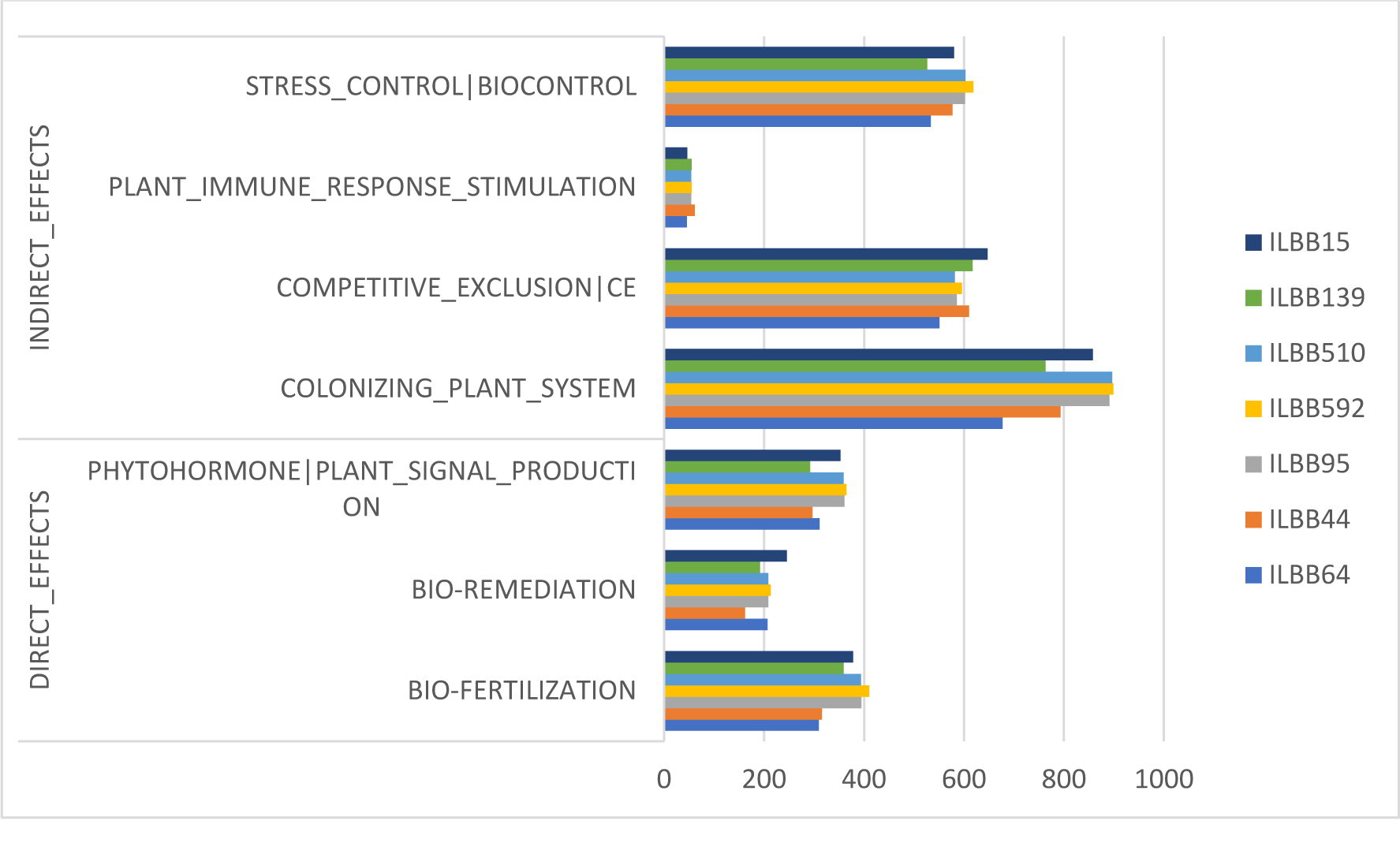
Number of genes present in each *Bacillus sensu lato* strain classified according to their direct and indirect effect on plant growth promotion.

**Figure 5.**
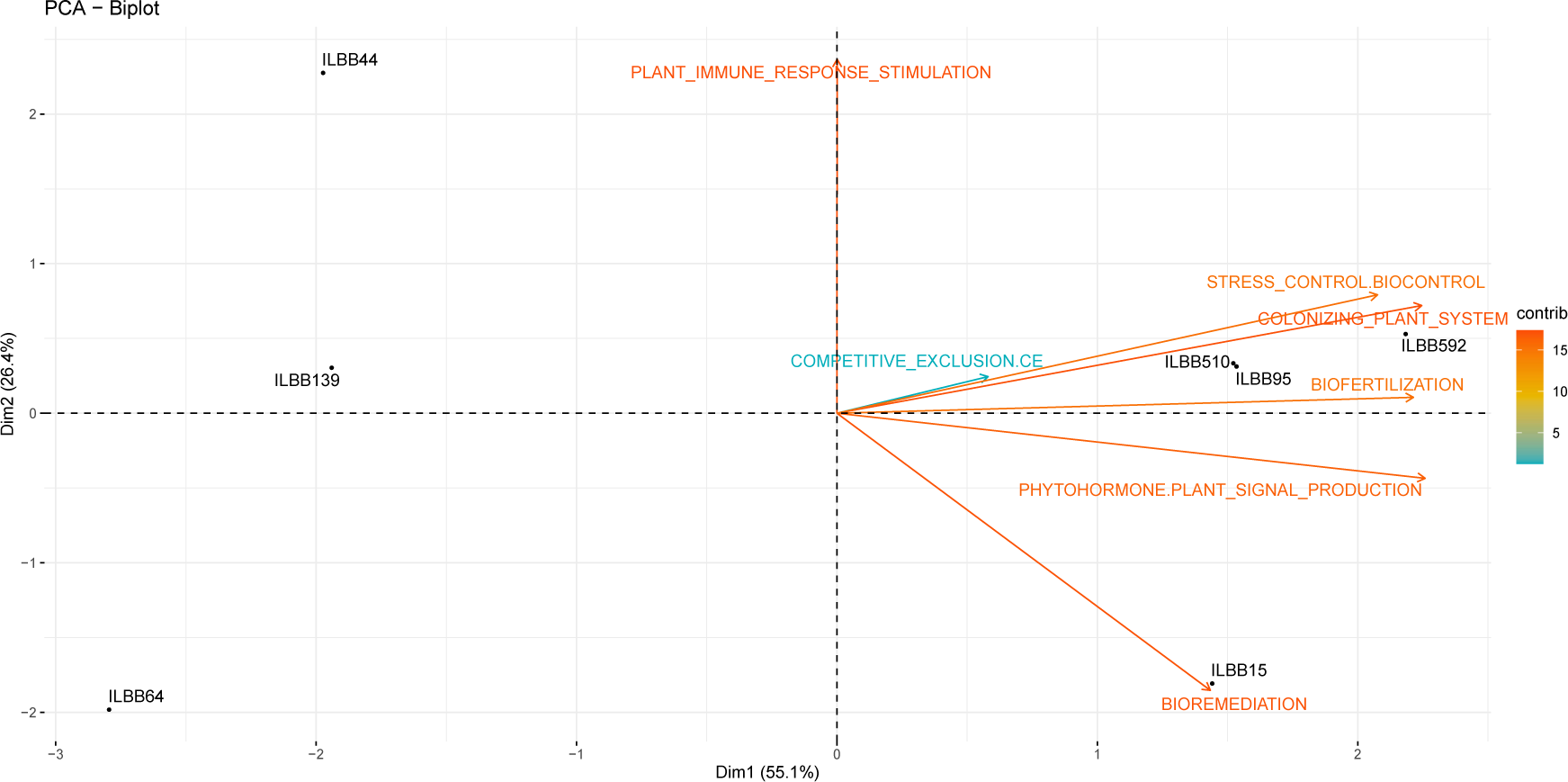
Ordination of *Bacillus sensu lato* strains by principal component analyses based on the presence/absence of genes involved in direct and indirect plant growth promotion.

Through genome mining using antiSMASH software version 5.0, 10 secondary metabolite gene clusters were identified in the genomes of the strain ILBB139, nine in ILBB44, eight in the strains ILBB15 and ILBB510, seven in ILBB64, six in the strains ILBB95 and ILBB592 (Supplementary Table 1, Supplementary Figure 1). All the strains encoded Terpene, NRPS-independent-siderophore and T3PKS, but only the strains ILBB15 and ILBB44 encoded betalactone, the strains ILBB15 and ILBB510 encoded lassopeptide, the strains ILBB95 and ILBB510 encoded lanthipeptide and the strains ILBB510 and ILBB592 encoded phophonate. Except ILBB139 and ILBB64, the strains showed clusters responsible for the synthesis of schizokinen (siderophore), while the synthesis surfactin was detected in the three *P. megaterium* strains and *B. pumilus* ILBB44. The compound citilin was detected only in the strain ILBB15. The correspondence analysis grouped the three *P. megaterium* strains based on the presence of BGC coding for carotenoid and surfactin, whereas the BGC for the lantibiotic PaenicidinA was present only in ILBB95 and ILBB510 (Fig. 6).

**Fig 6.**
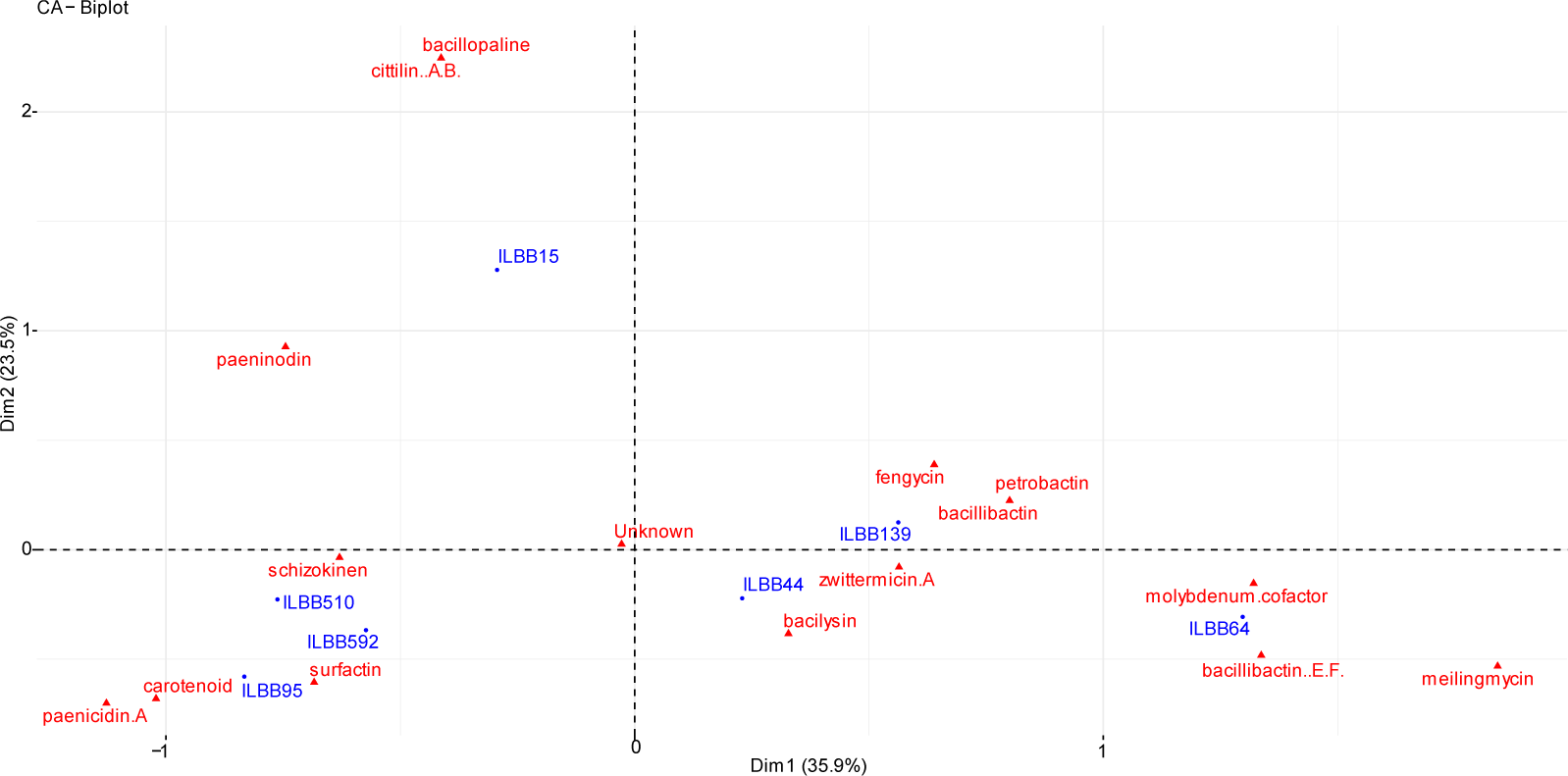
Ordination of *Bacillus sensu lato* strains by correspondence analyses based on the presence/absence of biosynthetic gene clusters.

The search for specific genes related with P mobilization showed that the three *P. megaterium* strains (ILBB592, 95 and 510), ILBB44 and ILBB15 were enriched in genes related with P solubilization and -except for ILBB44- also with genes involved in P mineralization from organic sources other than phytate. Regarding phytate degradation, sequences with homology to known phytases were found in all the strains. However, these sequences did not show the phytase domain. Regarding β-glucosidase genes involved in plant-rhizobia signalling for nodulation, ILBB44 had four and ILBB592 had three, whereas ILBB15 had one.

## Discussion

Soybean is one of the most extended crops and a fundamental protein source for human and animal consumption (Pagano & Miransari, 2016). Different from N, which is mostly sourced from atmospheric N through biological fixation by specialized rhizobia, P is added as fertilizer. However, there is an environmental need to reduce P additions while assuring satisfactory P and N nutrition of this crop. In this regard, immobilized forms of inorganic and organic P in soils could be targeted to improve their bioavailability to plants, and microbes with the capacity to solubilize and mineralize these unavailable P forms could therefore be developed as biological inputs or biofertilizers to achieve this goal (Mitter et al., 2021). Although plant phytases in soybeans are thought to be active in phytate mineralization, it is also acknowledged that organic substances in soybean root exudates can act as a substrate for microorganisms with additional phytase activity (Schwerdtner et al 2022). Therefore, phosphatase/phytase producing microorganisms, compatible with rhizobia and favoured by soybean root exudates, could play a role in P nutrition of soybeans. Members of *Bacillus sensu lato*, usually part of the rhizospheric community of plants, are known to produce organic acids, phosphatases and phytases and are considered convenient biological agents since their endospores can be easily produced and formulated for commercial use. In this work, a collection of *Bacillus sensu lato*, known to qualitatively solubilize and mineralize P *in vitro* were further evaluated for their capacity to quantitatively produce orthophosphate from inorganic and organic P forms. In addition, these strains were tested *in planta* and further characterized for the presence of traits related with plant growth promotion and rhizocompetence. Finally, their genomes were used to assign species -a key aspect for biofertilizers regulatory compliance- and analysed to explore the relationship between the observed phenotypes and the genetic makeup of the isolates.

Identification based on whole genome sequences performed with TYGS showed that ILBB95, ILBB592 and ILBB510 belonged to a clade including *Priestia megaterium* and *P. aryabhattai* type strains, in line with claims that the latter is a synonym of *P. megaterium* (Rao et al. 2019). Since ANIm values of these three strains with the type strain of *P. megaterium* were higher than with the type strain of *P. aryabhattai*, the three strains could be assigned to *P. megaterium sensu stricto*. Interestingly, the genome of ILBB44 showed higher identity with the pathogenic Bonn strain of *Bacillus pumilus* -originated from necrotic tissue in the hand of a boy (Grass et al. 2016)- than with the type strain of the species, both in the phylogenomic tree and regarding the ANIm value. This finding is two folds: it can point at the existence of a sister species of *B. pumilus* and, in the case of strain ILBB44, it alerts against its further development as a PGPR. ILBB139 was associated with the *Bacillus wiedmannii* type strain in the TYGS phylogram, although it was labelled as a different species. However, the ANIm value between these isolates was 96.43, above the usual threshold of 95-96% accepted for species delimitation (Chun et al. 2018), and therefore it was considered as a member of this species. The genome of ILBB64 formed a low support clade with *Lysinibacillus xylanilyticus* in the TYGS phylogenomic tree, and had an ANIm value with the type strain of the species of 91.63, below the 95-96 % threshold. Therefore, the species assignment for this isolate was inconclusive. Finally, ILBB15 was assigned to *Peribacillus butanolivorans* based on TYGS and ANIm values.

The initial *in vitro* assessment of bacterial mineralization and solubilization potential of the *Bacillus* strains allowed to rank *Pe. butanolivorans* (ILBB15), *Lysinybacillus* sp. (ILBB64) and *B. wiedmannii* (ILBB139) as the ones with highest P mineralization and P solubilization potential. On the lower level of potential mobilization *in vitro*, *B. pumilus s.l.* (ILBB 44) was followed by *Priestia megaterium* isolates, with two isolates characterized as positive for these traits (ILBB95, ILBB592) and the negative control isolate (ILBB510).

Due to the upmost importance of rhizobial nodulation for effective N fixation from the atmosphere, any *Bacillus* based P-biofertilizer needs to be compatible with rhizobia for their co-inoculation on soybean seeds. Therefore, we evaluated the effect of co-inoculation of the selected *Bacillus s.l.* with rhizobia on nodule biomass. Interestingly, only *P. megaterium* ILBB592 co-inoculated on seeds produced higher nodule dry weight than control plants, inoculated only with rhizobia, whereas *B. butanolivorans* ILBB15 showed the opposite effect, reducing the nodule biomass significantly. ILBB592 was positive for ACC deaminase and auxin production whereas ILBB15 was negative for both traits, which are known to positively influence nodulation (Alemneh et al. 2020; Flores Duarte et al. 2022). Both strains were also opposite to each other regarding the potential to form biofilm on soybean seed exudates: while exudates increased the biofilm forming potential of ILBB592, they reduced the biofilm capacity of ILBB15. Biofilm forming capacity is a key trait to assure bacterial colonization of the root, and can be stimulated by plant exudates (Beuregard et al. 2013). In this regard, it was notorious that ILBB592 outperformed all other strains when grown on soybean seeds exudates, whereas ILBB15 was the only strain negatively affected in its biofilm forming capacity by seed exudates. In addition, through genome mining, it was found that ILBB15 had fewer genes related with phytohormone plant signal, colonization of plant systems and biofertilization and more genes associated with bioremediation. On the contrary, those former PGP traits were highly represented in the genome of ILBB592 and the other two strains of the *P. megaterium*.

N nutrition expressed as N per plant -conditioned by N fixation at the nodules and N absorption from soil solution- was positively influenced by co-inoculation with ILBB592 in the sand peat vermiculite mix, together with ILBB139, ILBB44 and ILBB510, which had not shown a positive effect on nodule biomass. Because the nutritional evaluation was done only 60 days after sowing it is remarkable that plants inoculated with ILBB592 managed to increase both their nodulation and total plant N content, since it is known that supernodulation can produce compensating effects of decreased plant size and increased N concentration (Day et al. 1986), leading to variable outcomes in total N per plant.

When P nutrition was evaluated on two different substrate mixtures supplemented with phytate as organic P, *B. pumilus* ILBB44 and *P. megaterium* ILBB95 consistently produced higher P accumulation on a plant basis than control plants and most other isolates, while ILBB15 showed the lowest. The genome of ILBB44 was the smallest but it was characterized by the presence of genes related to stimulation of plant immune response, a category of genes in which it accumulated more genes than the other strains. However, because of the high ANIm value of this strain with pathogenic *B. pumilus* Bonn strain and the high number of genes associated with multidrug resistance, its development as a PGPR should not be pursued any further. On the contrary, ILBB95 belonging to *Priestia megaterium,* also enhanced P accumulation when grown on both substrate mixtures, although the effect was significantly different from control only on the poor mix without peat, when added phytate was the unique source of organic P. In the case of ILBB592, the higher nodulation stimulated by this strain was not mirrored by increased total P in aerial parts of the plant. However, this could be related to the higher demand for P exerted by the formation of the bigger nodular mass (Gunawardena et al., 1993). If this was the case, and because P was evaluated only on the aerial part of the plant, P consumed or accumulated in the higher nodular biomass was not evaluated and, therefore, it did not add to the total P per plant. In spite of this, the higher nodular biomass could be taken as a proxy for higher P bioavailability for the plant and rhizobia -and the nodulation process- when ILBB592 was co-inoculated with rhizobia.

The comparative analysis of the genomes, focused on direct and indirect plant growth promotion traits of the seven characterized strains, clearly separated the three *P. megaterium* strains from the others. The three *P. megaterium* strains were clearly associated with biofertilization, phytohormone plant signalling and plant colonization genetic determinants. The three strains of *P. megaterium* and ILBB15 of *Pe. butanolivorans* had higher numbers of genes related with inorganic P solubilization and organic P mineralization. In particular, *phoD* -which codes for an alkaline phosphatase with phosphomonoesterase activity and is considered the most relevant alkaline phosphatase gene in soil bacteria (Park et al 2022)- was present only in these four strains. Proteins with homology with phytases were found in all the strains but the known phytase domains were not present in any of them. In this regard, it would be necessary to verify the link between these sequences and the observed phenotypes. Similarly, Villamizar et al. (2019) found a gene product with phytase activity, lacking known catalytic domains associated with phytase activity, and suggested that it was a novel type of phytase.

In conclusion, the *in vitro* characterization comprising biofertilization, plant growth promotion, and rhizocompetence traits together with *in planta* assays and *in silico* genomic analysis of strains of *Bacillus sensu lato*, points at ILBB592 and ILBB95 -both ascribed to *Priestia megaterium*- as candidates to be used as PGPR with an impact on N and P nutrition of soybean plants. This species has already been associated with a positive effect on plant growth and also P solubilization (Kaminsky and Bell, 2022). Their development as biofertilizers with PGPR activity, alone or in a combined formulation, is envisaged.

## Acknowledgements

Agencia Nacional De Investigación e Innovación (ANII), Calister, Lage, Lafoner, and INIA for funding the projects. Personnel of Calister, Lage and Lafoner for their collaboration.

