## Supplementary material for "Phenotypic and genomic characterization of *Bacillus sensu lato* for their biofertilization effect and plant growth promotion features in soybean plants": suppl: figuras suplementarias.docx

Supplementary Table 1

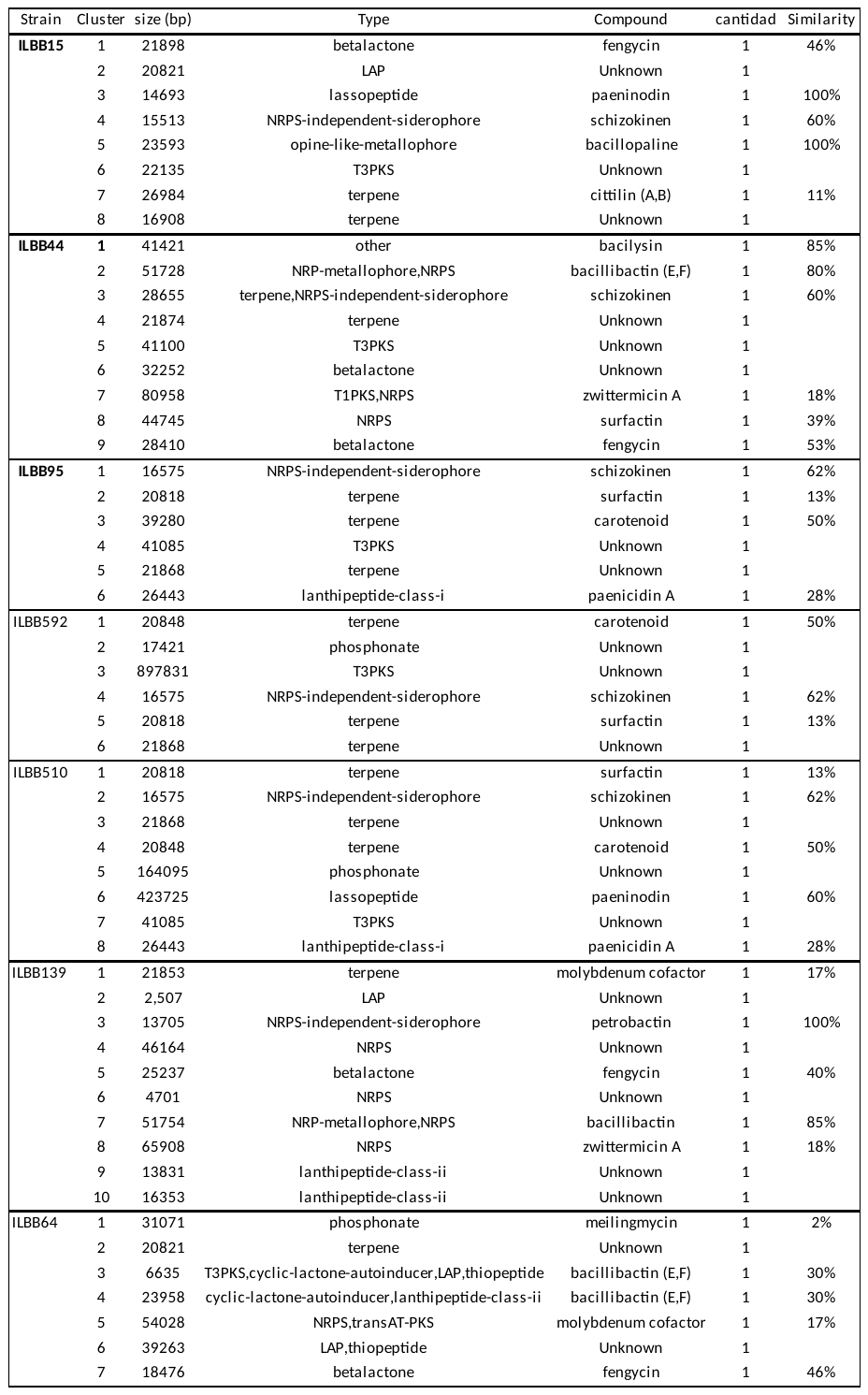

Supplementary Figure 1. Phylogenomic trees produced by TYGS.

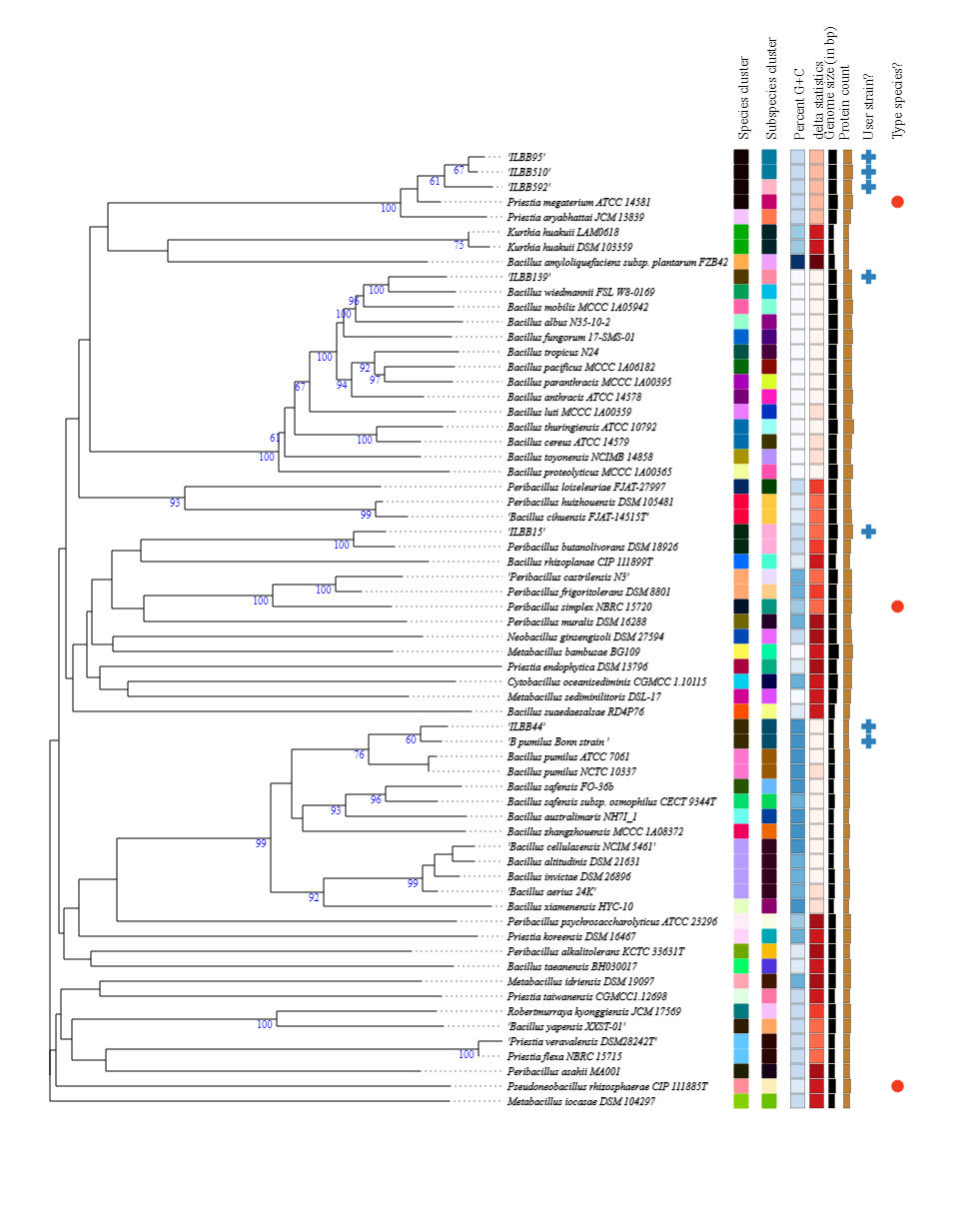

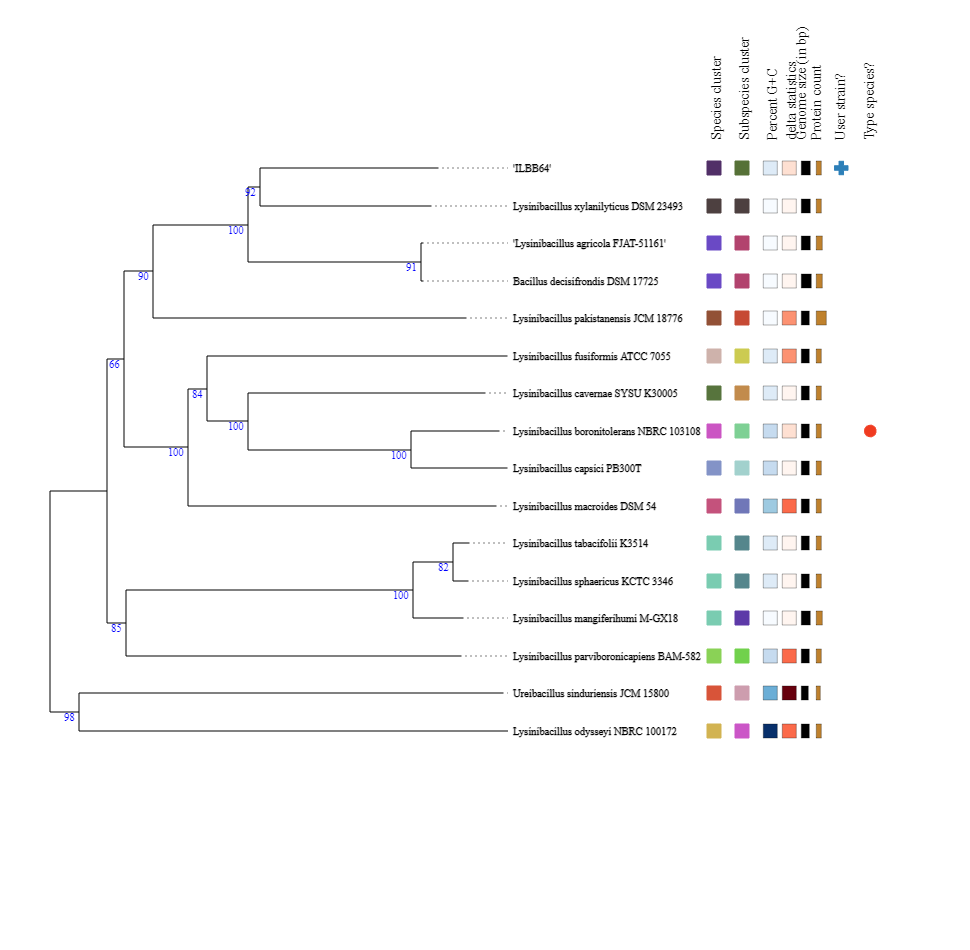

Supplementary Figure 2. Type and number of BGC present in each strain.

Genes clusters

| Strains | Unknown | schizokinen | surfactin | fengycin | carotenoid | bacillibactin (E,F) | paenicidin A | paeninodin | zwittermicin A | molybdenum cofactor | bacilysin | bacillopaline | cittilin (A,B) | petrobactin | bacillibactin | meilingmycin |
| --- | --- | --- | --- | --- | --- | --- | --- | --- | --- | --- | --- | --- | --- | --- | --- | --- |
| ILBB592 | 3 | 1 | 1 |  | 1 |  |  |  |  |  |  |  |  |  |  |  |
| ILBB95 | 2 | 1 | 1 |  | 1 |  | 1 |  |  |  |  |  |  |  |  |  |
| ILBB64 | 2 |  |  | 1 |  | 2 |  |  |  | 1 |  |  |  |  |  | 1 |
| ILBB510 | 3 | 1 | 1 |  | 1 |  | 1 | 1 |  |  |  |  |  |  |  |  |
| ILBB15 | 3 | 1 |  | 1 |  |  |  | 1 |  |  |  | 1 | 1 |  |  |  |
| ILBB44 | 3 | 1 | 1 | 1 |  | 1 |  |  | 1 |  | 1 |  |  |  |  |  |
| ILBB139 | 5 |  |  | 1 |  |  |  |  | 1 | 1 |  |  |  | 1 | 1 |  |
